# Study on The Effect of a Propiconazole Based Fungicide on Plant Growth Promoting Properties of Isolates from Wheat Rhizosphere

**DOI:** 10.64898/2026.10.21.760685

**Authors:** Mirza Tausif Haque, Ankita Saha, Anowara Begum, Humaira Akhter

**Affiliations:** Department of Microbiology, BRAC University, Dhaka, Bangladesh; Environmental Microbiology Laboratory, Department of Microbiology, University of Dhaka, Dhaka, Bangladesh

**Author notes:** **Correspondence:** Humaira Akhter.

**Keywords:** Rhizosphere, Propiconazole, PGPR, Variance, Correlation

## Abstract

Reliance on propiconazole based fungicides has been on the rise for better harvest of crops. Such fungicides can also affect non-target rhizosphere microorganisms. However, the off-target effect of propiconazole on rhizobacteria remains scarcely discussed. The main objective of the present study was to observe the effect of this compound on six plant growth promoting (PGP) properties of wheat rhizosphere isolates using a statistical approach. The study showed that although the overall PGP profile shifted only modestly across propiconazole doses, siderophore and ammonia production were suppressed at the highest fungicide concentration, IAA production declined in the presence of the fungicide, and cellulase activity declined at 4.24 mg/L and above, whereas siderophore appeared to act as a bridge between growth promotion and enzymatic activity. This pattern indicates propiconazole exerts a selective pressure on specific traits rather than holistic disruption. The findings of this study contribute to our understanding of pesticide risk assessment in sustainable agriculture.

## 1 Introduction

The rhizosphere is the narrow region of soil surrounding plant roots that is influenced by root exudates and associated microbial communities. Rhizobacteria that benefit their host, known as plant growth-promoting rhizobacteria (PGPR), promote plant growth by aiding promoting plant growth through aiding in nutrient uptake, disease resistance, stress tolerance, and enhancement of root development. These properties play a notable role in the development of cereal crops like rice, wheat, maize, oats, millet, making them a key element in sustainable agriculture.

Biofilms are structured communities of bacterial populations in the rhizosphereembedded in a matrix of extracellular polymeric substances (EPS) consisting of polysaccharides, proteins, extracellular DNA, and lipids (Haque *et al*., 2020). *Azotobacter, Pseudomonas, Serratia, Mesorhizobium* (Prasanna *et al*., 2011), *Bacillus subtilis, Azospirillum brasilense* (Ilyas *et al*., 2020), *Bacillus amyloliquefaciens* (Tan *et al*., 2013), *Rhizobium* (Rinaudi & Giordano, 2010), *Agrobacterium* (Ramey *et al*., 2004) are some of the biofilm producing bacteria often found in the rhizosphere population. According to Kasim *et al*. (2016), rhizosphere bacterial strains often resort to biofilm formation under varying stress conditions as a survival strategy. Inoculation of biofilm producing bacteria has been reported to increase drought tolerance (Karimi *et al*., 2022). PGPR mitigate salinity stress by producing exopolysaccharides, a critical component of bacterial biofilms (Li *et al*., 2021). Biofilm formation by *Pseudomonas stutzeri* A1501enables nitrogen fixation under aerobic conditions (Wang *et al*., 2017), and strains of *Bacillus subtilis* showed activity against *Ralstonia solanacearum*, a plant pathogen, which was dependent on widely conserved genes for biofilm formation (Chen *et al*., 2013). EPS, a component of biofilm, increases the water holding capacity of soil (Adessi *et al*., 2018), making biofilm a crucial element in sustainable agriculture.

Amylase is an extracellular enzyme catalyzing the hydrolysis of starch into sugars, playing a key role in the metabolism of carbohydrates in both plants and animals. *Bacillus cereus* (Huang *et al*., 2022a) and *Bacillus* sp. B51f (Srivathsan *et al*., 2022), *Enterobacter cloacae* (Luang-In *et al*., 2019) are examples of soil bacteria producing this enzyme. Amylase producing *Bacillus halotolerans* RFP74 has been shown to inhibit the growth of phytopathogens, which include *Alternaria* and *Bipolaris,* exceeding an inhibition rate of 60% (Rafanomezantsoa *et al*., 2023).

Cellulase, like amylase, is a common lytic enzyme in the rhizosphere. *Rhodococcus* sp., *Rhizobium* sp. (Silva *et al*., 2017), *Bacillus subtilis*, *B. mojavensis*, *B. cereus* (Balla *et al*., 2022), *Frankia*, *Azorhizobium, Bacillus, Paenibacillus, Brevibacillus*, *Pseudomonas* (Bhadrecha *et al*., 2020) are some of the bacterial members of rhizosphere microbiota capable of cellulase secretion. This enzyme plays a role in biocontrol activities against *Phytophthora capsici* (Admassie *et al*., 2022), and cellulase producing *Micromonospora carbonacea* showed biocontrol activity against *Phytophthora cinnamomi* (El-Tarabily *et al*., 1996). Application of exogenous cellulase increases soil urease and sucrase activities (Zhang *et al*., 2017) and has been reported to stimulate microbial growth (Xiao *et al*., 2024).

Siderophores are low-molecular-weight, iron-chelating compounds that help bacteria scavenge iron from their surroundings, thereby increasing the bioavailability of iron in soil. *Pantoea dispersa* MPJ9, *Pseudomonas putida* MPJ2 (Patel and Minocheherhomji, 2018), *Bacillus* MG214652 (Osman *et al*., 2018), *Streptomyces* sp. S29 (Jarmusch *et al*., 2021), *Rhizobium* (Manasa *et al*., 2017) are some of the examples of bacteria producing siderophores aiding plants in uptake of iron. Siderophore in soil has shown potential for plant growth and increased immunity (Trapet *et al*., 2016), enhancement of protein and carbohydrate content (Patel and Minocheherhomji, 2018), helping plants survive drought (Timofeeva *et al*., 2022), protection against fungal diseases (Jarmusch *et al*., 2021), mitigation of chlorosis (Ferreira *et al*., 2019), making these compounds an important part in plant-microbe interaction.

Among various phytohormones, Indole-3-acetic acid (IAA) is one of the most notable ones. It regulates plant growth and development, which includes the morphogenesis of plant shoots and roots along with their elongation (Naser & Shani, 2016). Along with this, IAA also helps in abiotic stress tolerance, including salt stress (Saini *et al*., 2021), sodic alkaline stress (Gong *et al*., 2014), and phosphorus-deficiency (Huang *et al*., 2022b). This phytohormone also scavenges reactive oxygen species, reduces Na^+^ accumulation, and protects photosystem II (PSII) from damage (Gong *et al*., 2014; Kaya *et al*., 2010). *Erwinia herbicola*, *Agrobacterium, Azospirillum, Bradyrhizobium, Enterobacter, Klebsiella, Pseudomonas,* and *Rhizobium* (Tsavkelova *et al*., 2006) are some examples of IAA producing bacteria in plant rhizosphere, indicating diversity in phytohormone producing rhizobacterial population.

*Serratia odorifera* 4Rx13, *S. plymuthica* HRO-C48, *S. plymuthica* 3Re4-18, *Pseudomonas fluorescens* L13-6-12, *P. fluorescens* 3Re2-7, *Bacillus subtilis* B2g, *Bacillus megaterium* PSB1, and *Paenibacillus polymyxa* COW3 are some of the bacterial species that emit high levels of ammonia in the rhizosphere (Weise *et al*., 2013). Ammonia released by PGPR can supply nitrogen to the host plant. However, high ammonia production can lead to alkalization of the rhizosphere, negatively impacting plant growth and development. For example, *Serratia odorifera* 4Rx13 and other rhizobacteria have been found to emit high levels of ammonia, causing alkalization of the plant medium and resulting in symptoms such as chlorosis of leaves, lowered root/shoot ratio, stimulated root branching, and inhibited seed germination and seedling establishment (Weise *et al*., 2013).

Application of pesticide in agricultural soil has advanced the world’s food production over the past decades. In a study conducted on 10 European countries and 8 cropping systems, total pesticide residue concentrations in samples were up to 28.7 mg kg^−1^ for conventionally managed fields and 5.46 mg kg^−1^ for organically managed fields (Knuth *et al*., 2024). Another study conducted in Gaidahawa Rural Municipality (GRM) (27° 35.429′ N and 83° 19.215’ E) of Nepal revealed the mean concentration of pesticides in soil samples was 16.05 μg kg^−1^ with a range of 1.02–251.28 μg kg^−1^ (Bhandari *et al.,* 2020). Maximum total residue concentration among the conventionally farmed soils was 3043 μg kg^−1^ during the sampling campaign of the Finnish national monitoring program for arable soils, conducted during the active growing season (June– August) in 2018 (Hagner *et al*., 2024).

Propiconazole is a systemic fungicide widely used in agriculture and horticulture. This fungicide belongs to the triazole class, acting as a demethylation inhibitor (DMI) disrupting the biosynthesis of ergosterol, a crucial component of fungal cell membranes. Besides fungi, propiconazole can affect soil bacterial populations too. Among soil bacterial communities, the relative abundance of *Proteobacteria* and *Planctomycetota* was reported to increase, while that of *Acidobacteria* decreased after propiconazole treatment (Sliti *et al.,* 2024). Propiconazole has also been shown to affect non-target bacterial endophyte diversity (Yadav *et al*., 2022), potentially compromising plant health. Several strains of *Pseudomonas putida* isolated from tea rhizosphere have been shown to degrade propiconazole *in vitro* (Sarkar *et al*., 2009).

In the current study, we aim to observe the effect of propiconazole based pesticide on six PGP properties (biofilm, amylase, cellulase, siderophore, IAA, ammonia) exerted by rhizosphere bacterial isolates and analyze the effect using a statistical approach. Globally, the topsoil in croplands has an average total pesticide content of 0.62 mg kg-soil^−1^ and a 95^th^ percentile of 2.12 mg kg-soil^−1^ (Tang & Maggi, 2021) - which was used as a benchmark for our test fungicide concentration. We emulated the concentration in laboratory culture medium at mg/L units and used double and triple the aforemnetioned value (4.24 mg/L and 6.36 mg/L, respectively) to observe effects of higher propiconazole concentrations.

## 2. Materials and Methods

### 2.1 Soil Sampling and Isolation of Bacteria

The study was performed on wheat rhizosphere soil samples collected from agronomy (24° 43′ 4.36″ N, 90° 25′ 31.43″ E) and entomology (24° 43′ 4.69″ N, 90° 25′ 31.58″ E) fields of Bangladesh Agricultural University, Mymensingh. Soil surrounding plant roots at about 20-25 cm depths were collected in labeled sterile zip lock bags, carried to the laboratory and were kept at 4°C for further study. One gram of soil sample was mixed with 10 mL of sterile 0.85% NaCl solution and the suspension was ten-fold serially diluted up to 10 dilution. From each dilution, 100 uL was spread on Yeast Extract Mannitol Agar (YMA) plates and were incubated at 30°C for 72 hours. Colonies were characterized morphologically based on their size, shape, color, elevation, margin, opacity and texture. A total of 30 isolates of different morphotypes were selected and streaked onto YMA media three consecutive times to obtain purified culture.

### 2.2 Assay of Plant Growth Promoting Properties

The effect of propiconazole was tested at four concentrations: 0 (untreated control), 2.12, 4.24 and 6.36 mg/L. Control cultures received no propiconazole. All other assay conditions were identical across the four concentrations.

#### 2.2.1 Biofilm Formation Assay

The ability of the rhizobacterial isolates to produce biofilms was assayed by the protocol described in O’Toole (2011) along with some modifications. LB broth was inoculated with one loopful of culture from a subculture plate and incubated at 30°C for 48 hours. The culture was then diluted hundredfold. 250 μL of the diluted culture was then introduced into a well in a transparent bottom microtiter plate in triplicate (three wells per isolate). A triplicate for negative control (containing only sterile broth) was also prepared. Three wells were left blank without any kind of inoculation which served as blank. The microtiter plate was incubated at 30°C for 24 hours and was washed after the incubation period. After drying, 125 μL 0.1% crystal violet was added to each well. After 10-15 minutes incubation at room temperature, the microtiter plate was washed again and dried. Afterwards, 125 μL 33% acetic acid was added to each well to solubilize the crystal violet. After another incubation period of 10-15 minutes at room temperature, the absorbance of crystal violet dissolved in acetic acid present in each well was observed at 575 nm wavelength using an ELISA reader. OD_cut_ was determined by the following equation:

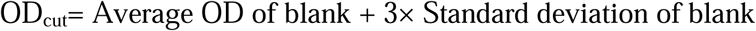

Degree of biofilm formation was determined using the following conditions:

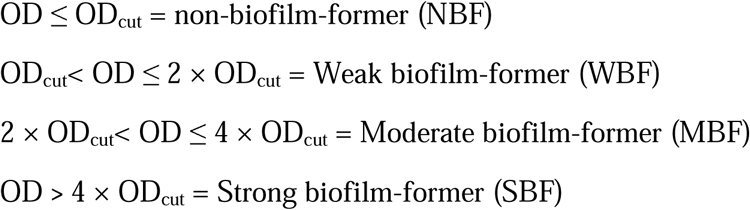

#### 2.2.2 Siderophore Production Assay

Universal Chrome Azurol S (CAS) (Schwyn and Neilands, 1987) assay was used to detect siderophore production ability of the isolates with some modifications. Bacterial culture was inoculated in 1 mL of modified siderophore broth [K_2_HPO_4_ 0.2 g; MgSO_4_ 0.2 g; Mannitol 10.0 g; (NH_4_)_2_SO_4_ 2 g; NaCl 0.05 g; PIPES free acid 32.24 g; Distilled water 1000.0 mL] and incubated at 30°C for 7 days. The culture was then centrifuged at 14000 rpm for 10 minutes. After that, 900 µL supernatant was added to 100 µL CAS reagent. 250 µL from the mixture was added to each microtiter well, using three wells per strain. The decrease in the intensity of blue color of the mixture within 20 minutes indicated siderophore production. An ELISA reader was used to measure the absorbances for each well at 630 nm. An uninoculated broth (treated with CAS reagent similar to the samples) was used as control. Siderophore index was then measured using this formula:

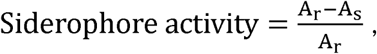

where A_r_ is the absorbance for sterile broth, and A_s_ is the absorbance for test sample. Any negative value for siderophore activity was interpreted as inability to produce siderophore.

#### 2.2.3 Indole-3-Acetic Acid (IAA) Production Assay

IAA production ability of the isolates was assessed by using LB broth amended with 0.5% L-tryptophan. 1mL LB broth with L-tryptophan was inoculated with the isolates and was incubated at 30°C for 72hrs (Shoukry *et al*., 2018). Uninoculated medium was incubated in parallel as a negative control. Following incubation, bacterial cells were removed by centrifugation at 4000 rpm for 20 minutes at 4°C. Then, 125 μL of cell free supernatant was introduced to each microtiter plate well, using three wells per isolate. Salkowski reagent [0.5M FeCl_3_ 15mL; Distilled water 500mL; H_2_SO_4_ (Concentrated) 300mL] was added in 1:1 ratio and incubated at room temperature for 20 minutes. The development of pink color indicated IAA production which was optically measured at 530 nm. Absorbance for Salkowski reagent added to the negative control media was deducted from each sample’s absorbance to omit any possible color development due to L-tryptophan. For the quantification of IAA production, a standard linear regression curve was prepared using different concentrations of IAA.IAA concentrations (µg/mL) were calculated from the regression equation.

#### 2.2.4 Ammonia Production Assay

Ammonia production by bacterial isolates was assessed by following the procedure mentioned in Abdelwahed *et al*. (2022) along with some volume adjustments. The bacterial cultures were inoculated in peptone broth and incubated at 30°C for 72 hours. After incubation, each culture was taken in an microcentrifuge tube and centrifuged at 14000 rpm for 10 minutes. 50 μL of supernatant from each tube was taken and added to 100 μL Nessler’s reagent. After a 10-minute incubation, 28 μL of the mixture was added to 165 μL distilled water in a microtiter well, giving a final volume of 193 µL. Each isolate was assayed in triplicate. After an incubation period of 10 minutes at room temperature, absorbance for each well was observed at 450 nm. Ammonia concentration was calculated by reference to ammonium sulfate standard range (μM) performed in the same conditions of the spectrophotometric assay.

#### 2.2.5 Cellulase Production Assay

The cellulase activity of the rhizobacterial isolates under different propiconazole treatments was measured using the 3,5-dinitrosalicylic acid (DNS) method (Deshavath *et al*., 2020; Pan, 2017). To extract the crude enzyme, each isolate was inoculated in CMC broth [Peptone 5.0 g; NaCl 5.0 g; Yeast extract 3.0 g; Carboxymethyl cellulose 5.0 g; Distilled water 1000 mL]. Four concentrations of propiconazole (0 mg/L, 2.12 mg/L, 4.24 mg/L, and 6.36 mg/L) were added, and the cultures were incubated at 30°C and 150 rpm for 2 days. The culture broth from each treatment was aseptically collected and diluted twofold with fresh medium before the assay. This diluted culture was used as the crude enzyme source. A volume of 0.5 mL of the prepared enzyme solution from each treatment was added to a stoppered test tube containing 1.5 mL of 1% (w/v) sodium carboxymethyl cellulose (CMC-Na) solution. The mixture was incubated in a water bath at 50°C for 30 minutes. The reaction was terminated by the addition of 1.5 mL of DNS reagent. The tubes were then immersed in a boiling water bath for 5 minutes. After cooling, the reaction volume was adjusted to 25 mL with distilled water, and the absorbance was recorded at 540 nm using an ELISA reader. Cellulase activity was calculated using the equation:

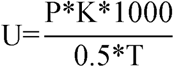

where U is the enzyme activity, P is the glucose content from the standard curve, K is the dilution factor of the enzyme solution, and T is the incubation time (min) in the constant-temperature water bath.

For the glucose standard curve, a working stock of 1.0 mg/mL glucose was used to prepare six graduated, stoppered test tubes. These were prepared with 0.0, 0.4, 0.8, 1.2, 1.6, and 2.0 mL of the glucose standard, and distilled water was added in inverse proportion (2.0, 1.6, 1.2, 0.8, 0.4, and 0.0 mL) ‘to keep the total volume constant. Then, 1.5 mL of DNS reagent was added to each tube, followed by thorough mixing and boiling for 5 minutes. After cooling to room temperature under running water, the volume was adjusted with distilled water, and the solutions were mixed again before the absorbance was recorded at 540 nm.

A linear regression equation was established with glucose concentration as the independent variable and absorbance as the dependent variable. This equation was used to calculate the amount of glucose (P) released in each enzymatic reaction across all four propiconazole treatments.

#### 2.2.6 Amylase Production Assay

The amylase activity of the rhizobacterial isolates under different propiconazole treatments was measured using the 3,5-dinitrosalicylic acid (DNS) method (Salihu *et al*., 2015). For crude enzyme extraction, each isolate was inoculated into Luria Bertani broth with four different concentrations of propiconazole: 0 mg/L, 2.12 mg/L, 4.24 mg/L, and 6.36 mg/L. The cultures were incubated at 30°C for 2 days. After incubation, the culture broth from each treatment was centrifuged at 10,000 rpm for 15 minutes at 4°C. The cell-free supernatant was used as the crude enzyme source. The reaction mixture was prepared by combining 250 µL of the crude enzyme extract with 250 µL of a 1% soluble starch solution, prepared in 0.2 M phosphate buffer, pH 7.0. This mixture was incubated at 37°C for 10 minutes. The reaction was stopped by the addition of 500 µL DNS reagent, followed by boiling in a water bath for 5 minutes and rapid cooling under running tap water. The mixture was then diluted with 1 mL deionised distilled water, and the absorbance was measured at 540 nm using a spectrophotometer or ELISA reader. A blank was also prepared. The amount of reducing sugar produced was measured against a standard curve created under the same assay conditions.

A glucose standard curve was created using a working stock solution of 1.0 mg/mL glucose. Six graduated, stoppered test tubes were prepared, and the glucose standard solution was added to them at volumes of 0.0, 0.4, 0.8, 1.2, 1.6, and 2.0 mL Distilled water was added to each tube in inverse proportion to the glucose volume (2.0, 1.6, 1.2, 0.8, 0.4, and 0.0 mL, respectively) to maintain a constant total volume across all tubes. For color development, the same reagent and method used in the amylase assay were applied to each glucose standard. Specifically, 500 µL of DNS reagent was added to each standard, followed by heating in a boiling water bath to develop the color. After cooling to room temperature, the absorbance of each solution was measured at 540 nm using a spectrophotometer. The blank, in which distilled water was used instead of glucose, was used for zero adjustment.

## 3 Results

### 3.1 Biofilm Formation Assay

All 30 isolates were tested for their biofilm formation ability under 4 different propiconazole concentrations (0 mg/L, 2.12 mg/L, 4.24 mg/L, and 6.36 mg/L). All isolates except isolate 56 formed biofilm to some extent under at least one treatment of pesticide. In all doses, most of the isolates were weak biofilm formers (19 in 0 mg/L, 16 in 2.12 mg/L, 18 in 4.24 mg/L, and 18 in 6.36 mg/L) (Table 1). Twelve isolates showed weak biofilm production under all treatments, whereas three showed strong levels of biofilm in all four propiconazole concentrations - indicating that biofilm formation in half of the isolates was unaffected by the fungicide.

**Table 1:** Degree of biofilm formation by isolates under different pesticide concentrations.

| Isolate | 0 mg/L | 2.12 mg/L | 4.24 mg/L | 6.36 mg/L |
| --- | --- | --- | --- | --- |
| 22 | Moderate | Strong | Moderate | Strong |
| 23 | None | Weak | Strong | None |
| 24 | Weak | Weak | Weak | Weak |
| 26 | Strong | Strong | Strong | Strong |
| 27 | Weak | Weak | Weak | Weak |
| 29 | Weak | Weak | Strong | Strong |
| 30 | Weak | Weak | Weak | Weak |
| 34 | Weak | Weak | Weak | Weak |
| 35 | Weak | Moderate | Weak | Weak |
| 36 | Weak | Weak | Weak | Weak |
| 38 | Weak | Weak | Weak | Weak |
| 39 | Weak | Weak | Weak | Weak |
| 40 | Weak | Weak | Weak | Weak |
| 42 | Weak | Moderate | Moderate | Weak |
| 44 | Weak | Weak | Weak | Weak |
| 45 | Weak | Weak | Weak | Weak |
| 46 | Weak | Weak | Weak | Weak |
| 47 | Weak | Weak | Weak | Weak |
| 48 | Strong | Strong | Moderate | Moderate |
| 52 | None | None | Weak | Weak |
| 53 | Strong | Strong | Strong | Strong |
| 54 | Weak | None | Weak | Weak |
| 55 | Strong | Strong | Strong | Strong |
| 56 | None | None | None | None |
| 57 | None | Weak | Weak | Strong |
| 58 | Weak | None | Weak | Weak |
| 59 | Weak | None | None | None |
| 60 | None | None | None | Weak |
| 63 | Strong | Weak | Moderate | Moderate |
| 65 | Weak | None | Weak | None |

For statistical analysis, biofilm categories were coded as ordinal scores (none = 0, weak = 1, moderate = 2, strong = 3; Table 2).

**Table 2:** Numerical values assigned to degree of biofilm formation.

| Isolate | 0 mg/L | 2.12 mg/L | 4.24 mg/L | 6.36 mg/L |
| --- | --- | --- | --- | --- |
| 22 | 2 | 3 | 2 | 3 |
| 23 | 0 | 1 | 3 | 0 |
| 24 | 1 | 1 | 1 | 1 |
| 26 | 3 | 3 | 3 | 3 |
| 27 | 1 | 1 | 1 | 1 |
| 29 | 1 | 1 | 3 | 3 |
| 30 | 1 | 1 | 1 | 1 |
| 34 | 1 | 1 | 1 | 1 |
| 35 | 1 | 2 | 1 | 1 |
| 36 | 1 | 1 | 1 | 1 |
| 38 | 1 | 1 | 1 | 1 |
| 39 | 1 | 1 | 1 | 1 |
| 40 | 1 | 1 | 1 | 1 |
| 42 | 1 | 2 | 2 | 1 |
| 44 | 1 | 1 | 1 | 1 |
| 45 | 1 | 1 | 1 | 1 |
| 46 | 1 | 1 | 1 | 1 |
| 47 | 1 | 1 | 1 | 1 |
| 48 | 3 | 3 | 2 | 2 |
| 52 | 0 | 0 | 1 | 1 |
| 53 | 3 | 3 | 3 | 3 |
| 54 | 1 | 0 | 1 | 1 |
| 55 | 3 | 3 | 3 | 3 |
| 56 | 0 | 0 | 0 | 0 |
| 57 | 0 | 1 | 1 | 3 |
| 58 | 1 | 0 | 1 | 1 |
| 59 | 1 | 0 | 0 | 0 |
| 60 | 0 | 0 | 0 | 1 |
| 63 | 3 | 1 | 2 | 2 |
| 65 | 1 | 0 | 1 | 0 |

### 3.2 Siderophore Production Assay

The Siderophore indices for all isolates were calculated under all four treatments of the fungicide (Table 3). Four isolates (29, 40, 48, and 54) produced no siderophore in any of the pesticide doses.

**Table 3:** Siderophore indices of isolates under different pesticide concentrations.

| Isolate | 0 mg/L | 2.12 mg/L | 4.24 mg/L | 6.36 mg/L | Average Siderophore Index |
| --- | --- | --- | --- | --- | --- |
| 22 | 0.256 | 0.358 | 0.36 | 0 | 0.244 |
| 23 | 0.227 | 0.852 | 0.883 | 0.22 | 0.546 |
| 24 | 0.802 | 0.937 | 0.939 | 0.661 | 0.835 |
| 26 | 0.929 | 0.928 | 0.258 | 0.86 | 0.744 |
| 27 | 0.113 | 0.168 | 0.182 | 0 | 0.116 |
| 29 | 0 | 0 | 0 | 0 | 0.000 |
| 30 | 0.257 | 0.545 | 0.527 | 0.004 | 0.333 |
| 34 | 0.142 | 0.217 | 0.246 | 0 | 0.151 |
| 35 | 0.275 | 0.354 | 0.281 | 0.277 | 0.297 |
| 36 | 0.209 | 0.318 | 0.286 | 0.386 | 0.300 |
| 38 | 0.344 | 0.362 | 0.319 | 0.052 | 0.269 |
| 39 | 0.222 | 0.393 | 0.362 | 0.081 | 0.265 |
| 40 | 0 | 0 | 0 | 0 | 0.000 |
| 42 | 0.151 | 0.058 | 0.039 | 0 | 0.062 |
| 44 | 0.318 | 0.282 | 0.333 | 0.072 | 0.251 |
| 45 | 0 | 0.019 | 0 | 0 | 0.005 |
| 46 | 0.101 | 0.177 | 0.361 | 0.065 | 0.176 |
| 47 | 0.081 | 0.152 | 0.1 | 0 | 0.083 |
| 48 | 0 | 0 | 0 | 0 | 0.000 |
| 52 | 0.013 | 0 | 0 | 0 | 0.003 |
| 53 | 0.365 | 0.553 | 0.763 | 0.134 | 0.454 |
| 54 | 0 | 0 | 0 | 0 | 0.000 |
| 55 | 0.43 | 0.04 | 0.161 | 0.271 | 0.226 |
| 56 | 0.354 | 0.287 | 0.206 | 0.269 | 0.279 |
| 57 | 0.445 | 0.433 | 0.431 | 0.246 | 0.389 |
| 58 | 0.386 | 0.302 | 0.285 | 0.205 | 0.295 |
| 59 | 0.01 | 0 | 0 | 0 | 0.003 |
| 60 | 0 | 0.294 | 0.428 | 0 | 0.181 |
| 63 | 0.117 | 0.061 | 0.1 | 0.19 | 0.117 |
| 65 | 0 | 0 | 0 | 0.004 | 0.001 |
| Average | 0.218 | 0.270 | 0.262 | 0.133 |  |

### 3.3 IAA Production Assay

Out of 30 isolates, 19 achieved the highest IAA concentration in the absence of fungicide. Isolate 29 reached its peak IAA level at 2.12 mg/L propiconazole. Meanwhile, 5 isolates showed maximum production at 4.24 mg/L, and 5 isolates at 6.36 mg/L.

**Table 4:** IAA production (μg/mL) by isolates under different pesticide concentrations.

| Isolate | 0 mg/L | 2.12 mg/L | 4.24 mg/L | 6.36 mg/L | Average IAA Production |
| --- | --- | --- | --- | --- | --- |
| 22 | 3.18 | 6.957 | 4.524 | 8.066 | 5.682 |
| 23 | 45.096 | 20.688 | 22.262 | 33.251 | 30.324 |
| 24 | 31.992 | 51.775 | 68.056 | 76.008 | 56.958 |
| 26 | 61.916 | 52.21 | 54.206 | 53.745 | 55.519 |
| 27 | 94.521 | 80.543 | 54.762 | 61.77 | 72.899 |
| 29 | 17.586 | 20.254 | 17.183 | 14.856 | 17.470 |
| 30 | 12.452 | 6.739 | 0.675 | 5.062 | 6.232 |
| 34 | 12.031 | 1.232 | 0 | 1.646 | 3.727 |
| 35 | 12.107 | 3.442 | 0 | 1.975 | 4.381 |
| 36 | 13.908 | 0.507 | 0.079 | 2.84 | 4.334 |
| 38 | 0.69 | 0.652 | 2.54 | 2.222 | 1.526 |
| 39 | 12.874 | 4.275 | 0 | 0 | 4.287 |
| 40 | 6.82 | 3.116 | 0 | 0 | 2.484 |
| 42 | 17.126 | 14.529 | 13.849 | 10.576 | 14.020 |
| 44 | 5.249 | 1.486 | 0 | 0 | 1.684 |
| 45 | 5.249 | 1.775 | 0 | 0.7 | 1.931 |
| 46 | 12.95 | 0 | 0.873 | 0 | 3.456 |
| 47 | 10.996 | 5.833 | 1.587 | 0.947 | 4.841 |
| 48 | 60.421 | 51.594 | 38.175 | 37.243 | 46.858 |
| 52 | 0.852 | 1.379 | 3.59 | 0.891 | 1.678 |
| 53 | 66.185 | 52.682 | 66.197 | 59.574 | 61.160 |
| 54 | 36.148 | 16.935 | 34.829 | 38.333 | 31.561 |
| 55 | 75.852 | 63.065 | 59.744 | 57.752 | 64.103 |
| 56 | 47.481 | 41.418 | 58.376 | 50.969 | 49.561 |
| 57 | 56.593 | 46.245 | 38.077 | 51.279 | 48.049 |
| 58 | 70.741 | 57.625 | 55.256 | 57.752 | 60.344 |
| 59 | 74.185 | 44.636 | 38.825 | 57.829 | 53.869 |
| 60 | 2.111 | 0 | 2.179 | 0 | 1.073 |
| 63 | 5.037 | 1.073 | 4.957 | 14.496 | 6.391 |
| 65 | 3.778 | 5.134 | 5.385 | 8.45 | 5.687 |
| Average | 29.204 | 21.927 | 21.540 | 23.608 |  |

### 3.4 Ammonia Production Assay

All isolates in the present study were capable of ammonia production under all propiconazole treatments. Isolate 40 had the lowest average ammonia concentration and isolate 44 had the highest. Average ammonia production ranged between 2500–5800 μM (Table 5). Ten isolates (22, 27, 35, 36, 39, 52, 54, 58, 60 and 63) produced the highest concentration of ammonia when not challenged with propiconazole, which makes up one-third of the study population. The number was 11 in case of 2.12 mg/L propiconazole. Six isolates had maximum ammonia concentration in 4.24 mg/L fungicide treatment, and only three had the maximum in 6.36 mg/L.

**Table 5:** Ammonia production (μM) by isolates under different pesticide concentrations.

| Isolate | 0 mg/L | 2.12 mg/L | 4.24 mg/L | 6.36 mg/L | Average Ammonia Production |
| --- | --- | --- | --- | --- | --- |
| 22 | 8581.99 | 3361.46 | 4682.69 | 3848 | 5118.54 |
| 23 | 2621.81 | 5330.39 | 3706.63 | 2446.69 | 3526.38 |
| 24 | 3587.87 | 4541.14 | 2288.58 | 2911.8 | 3332.35 |
| 26 | 2642.81 | 2561.71 | 1903.89 | 4547.66 | 2914.02 |
| 27 | 5423.39 | 4793.03 | 3416.06 | 2067.04 | 3924.88 |
| 29 | 3419.86 | 5099.5 | 3927.62 | 3251.7 | 3924.67 |
| 30 | 4591.73 | 5235.94 | 2145.34 | 2357.24 | 3582.56 |
| 34 | 4289.31 | 4887.49 | 3729.14 | 866.48 | 3443.11 |
| 35 | 5709.01 | 4058.35 | 4324.6 | 2122.7 | 4053.67 |
| 36 | 5490.59 | 1999.16 | 4699.06 | 2681.23 | 3717.51 |
| 38 | 4814.35 | 5269.52 | 5259.73 | 2216.12 | 4389.93 |
| 39 | 4994.96 | 3890.43 | 2026.66 | 2587.81 | 3374.97 |
| 40 | 1336.53 | 4895.89 | 2034.85 | 1735.1 | 2500.59 |
| 42 | 576.28 | 5494.12 | 1918.21 | 8463.38 | 4113.00 |
| 44 | 4276.71 | 8287.99 | 5834.72 | 4732.51 | 5782.98 |
| 45 | 1021.51 | 3896.73 | 2726.48 | 3223.87 | 2717.15 |
| 46 | 4482.53 | 5439.55 | 3105.03 | 2416.87 | 3861.00 |
| 47 | 3377.86 | 4866.5 | 5341.58 | 2206.18 | 3948.03 |
| 48 | 5188.83 | 4827.79 | 7016.23 | 4924.67 | 5489.38 |
| 52 | 5433.56 | 5129.6 | 5309.78 | 1991.57 | 4466.13 |
| 53 | 4688.25 | 3400.8 | 6785.91 | 3472.62 | 4586.90 |
| 54 | 5589.3 | 5478.03 | 5519.16 | 3741.38 | 5081.97 |
| 55 | 5945.27 | 6130.49 | 7832.81 | 2387.94 | 5574.13 |
| 56 | 2513.49 | 4253 | 5963.04 | 4884.07 | 4403.40 |
| 57 | 3180.93 | 6640.92 | 4413.63 | 3027.92 | 4315.85 |
| 58 | 7063.24 | 3953.4 | 5975.61 | 4251.82 | 5311.02 |
| 59 | 5155.46 | 4881.05 | 5309.78 | 2358.93 | 4426.31 |
| 60 | 6568.22 | 2364.4 | 3923.68 | 1858.16 | 3678.62 |
| 63 | 5183.8 | 2564.14 | 4547.63 | 3327.61 | 3905.80 |
| 65 | 2646.13 | 2945.85 | 3848.3 | 4673.32 | 3528.40 |
| Average | 4346.52 | 4549.28 | 4317.21 | 3186.08 |  |

### 3.5 Cellulase Production Assay

Ten isolates had cellulase activity in all four treatments whereas four isolates produced no cellulase (Table 6).

**Table 6:** Cellulase activity (µg glucose/min/mL) by isolates under different pesticide concentrations.

| Isolate | 0 mg/L | 2.12 mg/L | 4.24 mg/L | 6.36 mg/L | Average Cellulase Activity |
| --- | --- | --- | --- | --- | --- |
| 22 | 0.000 | 0.000 | 0.000 | 0.000 | 0.000 |
| 23 | 0.000 | 42.608 | 3.213 | 12.536 | 14.589 |
| 24 | 64.994 | 46.876 | 44.868 | 50.100 | 51.710 |
| 26 | 0.000 | 0.000 | 0.000 | 0.000 | 0.000 |
| 27 | 0.000 | 52.309 | 34.157 | 19.289 | 26.439 |
| 29 | 0.000 | 0.000 | 0.000 | 0.000 | 0.000 |
| 30 | 74.496 | 48.428 | 14.321 | 48.834 | 46.520 |
| 34 | 88.940 | 50.369 | 5.990 | 0.000 | 36.325 |
| 35 | 109.464 | 83.741 | 0.000 | 20.977 | 53.545 |
| 36 | 80.578 | 67.831 | 37.727 | 26.886 | 53.255 |
| 38 | 60.053 | 27.086 | 9.561 | 0.000 | 24.175 |
| 39 | 116.306 | 53.085 | 54.785 | 45.879 | 67.514 |
| 40 | 50.931 | 32.906 | 30.983 | 8.315 | 30.784 |
| 42 | 46.370 | 40.667 | 3.213 | 23.932 | 28.546 |
| 44 | 82.098 | 52.697 | 53.595 | 31.529 | 54.980 |
| 45 | 57.013 | 44.936 | 13.924 | 24.776 | 35.162 |
| 46 | 73.736 | 65.503 | 0.000 | 8.315 | 36.888 |
| 47 | 97.681 | 24.757 | 2.420 | 40.815 | 41.418 |
| 48 | 121.627 | 30.190 | 0.000 | 0.000 | 37.954 |
| 52 | 0.000 | 0.000 | 0.000 | 0.000 | 0.000 |
| 53 | 0.000 | 16.997 | 6.387 | 27.308 | 12.673 |
| 54 | 42.569 | 34.847 | 0.000 | 0.000 | 19.354 |
| 55 | 0.000 | 11.952 | 2.420 | 19.289 | 8.415 |
| 56 | 0.000 | 25.534 | 42.091 | 35.328 | 25.738 |
| 57 | 0.000 | 36.787 | 6.784 | 27.730 | 17.825 |
| 58 | 72.976 | 34.071 | 0.000 | 44.191 | 37.809 |
| 59 | 55.872 | 46.488 | 0.436 | 0.000 | 25.699 |
| 60 | 0.000 | 15.832 | 29.793 | 45.457 | 22.771 |
| 63 | 0.000 | 2.251 | 0.000 | 0.000 | 0.563 |
| 65 | 86.279 | 49.205 | 37.727 | 33.217 | 51.607 |
| Average | 46.066 | 34.598 | 14.480 | 19.823 |  |

### 3.6 Amylase Production Assay

Nine isolates showed no amylase activity under any of the four treatments (Table 7). On the other hand, isolates 39 and 65 retained their amylase activity under all treatments.

**Table 7:** Amylase activity (µg glucose/min/mL) by isolates under different pesticide concentrations.

| Isolate | 0 mg/L | 2.12 mg/L | 4.24 mg/L | 6.36 mg/L | Average Amylase Activity |
| --- | --- | --- | --- | --- | --- |
| 22 | 0.000 | 0.000 | 0.000 | 0.000 | 0.000 |
| 23 | 2.664 | 0.000 | 0.000 | 0.000 | 0.666 |
| 24 | 0.000 | 0.000 | 151.927 | 136.515 | 72.110 |
| 26 | 0.000 | 0.000 | 0.000 | 0.000 | 0.000 |
| 27 | 16.516 | 0.000 | 0.000 | 0.000 | 4.129 |
| 29 | 5.860 | 0.000 | 0.000 | 0.000 | 1.465 |
| 30 | 96.430 | 27.240 | 0.000 | 59.906 | 45.894 |
| 34 | 119.872 | 17.315 | 51.898 | 0.000 | 47.271 |
| 35 | 3.729 | 0.000 | 0.000 | 0.879 | 1.152 |
| 36 | 18.647 | 11.800 | 0.000 | 0.000 | 7.612 |
| 38 | 4.795 | 0.000 | 0.000 | 13.438 | 4.558 |
| 39 | 14.385 | 49.297 | 50.721 | 52.370 | 41.693 |
| 40 | 0.000 | 83.485 | 51.898 | 0.000 | 33.846 |
| 42 | 0.000 | 0.000 | 0.000 | 0.000 | 0.000 |
| 44 | 34.630 | 23.932 | 0.000 | 3.391 | 15.488 |
| 45 | 0.000 | 27.240 | 0.000 | 0.000 | 6.810 |
| 46 | 0.000 | 0.000 | 0.000 | 0.000 | 0.000 |
| 47 | 0.000 | 15.109 | 0.000 | 0.000 | 3.777 |
| 48 | 3.729 | 0.000 | 0.000 | 0.000 | 0.932 |
| 52 | 0.000 | 0.000 | 0.000 | 0.000 | 0.000 |
| 53 | 17.581 | 0.000 | 0.000 | 0.000 | 4.395 |
| 54 | 2.664 | 0.000 | 0.000 | 0.000 | 0.666 |
| 55 | 0.000 | 0.000 | 0.000 | 0.000 | 0.000 |
| 56 | 0.533 | 28.343 | 0.000 | 3.391 | 8.067 |
| 57 | 0.000 | 0.000 | 0.000 | 0.000 | 0.000 |
| 58 | 4.795 | 0.000 | 0.000 | 0.000 | 1.199 |
| 59 | 0.000 | 0.000 | 0.000 | 0.000 | 0.000 |
| 60 | 5.860 | 33.857 | 0.000 | 0.000 | 9.929 |
| 63 | 0.000 | 0.000 | 0.000 | 0.000 | 0.000 |
| 65 | 18.647 | 11.800 | 10.709 | 46.091 | 21.812 |
| Average | 12.378 | 10.981 | 10.572 | 10.533 |  |

### 3.7 Statistical Analysis

Several statistical methods were applied to check the variance present in PGP traits, and to find out the interrelations between them. These tests include isolate-blocked permutation tests, multivariate PERMANOVA, PERMDISP, Wilcoxon signed-rank test, Kruskal–Wallis test, principal component analysis (PCA) with Friedman tests on component scores, redundancy analysis (RDA), and ordinal logistic regression. All the statistical tests were conducted using Python scripts on Google Colab platform. Because the same 30 isolates were assayed at all four propiconazole concentrations, the data were treated as repeated measures wherever the test allowed.

#### 3.7.1 Normality Test

Shapiro–Wilk and Kolmogorov–Smirnov tests were used to determine whether the PGPs were normally distributed or not in order to find out the applicability of a statistical test. Siderophore, IAA, biofilm, and amylase data were not normally distributed under any pesticide treatment (Supplementary Table 1, 2, 4, and 6). On the other hand, ammonia production was normally distributed except under the highest fungicide dose (Supplementary Table 3), and cellulase production followed normal distribution only in the 2.12 mg/L pesticide dose (Supplementary Table 5).

#### 3.7.2 Wilcoxon Signed-Rank Test

Since most of the data were not normally distributed, Wilcoxon signed-rank test was appliedto compare each pair of concentrations within isolates. Biofilm and amylase production appeared to have no remarkable differences across different propiconazole doses (Supplementary Tables 10 and 12). Mean siderophore production and mean ammonia production by bacterial population was lower than any other doses in case of 6.36 mg/L pesticide treatment (Supplementary Tables 7 and 9), and mean IAA production was significantly higher when the population faced no fungicide (Supplementary Table 8). Cellulase production was effectively similar under 0 mg/L and 2.12 mg/L propiconazole doses, which is also true for 4.24 mg/L and 6.36 mg/L treatments (Supplementary Table 11).

#### 3.7.3 Isolate-Blocked Permutation Test and Multivariate PERMANOVA

##### 3.7.3.1 Isolate-Blocked Permutation Test

Dose effects on each trait were tested with an isolate-blocked (repeated-measures) permutation test, in which concentration labels were permuted within each isolate (4,999 permutations). Global p-values were Holm-adjusted across the six traits, and pairwise p-values were adjusted using the Benjamini–Hochberg false discovery rate (FDR) within each trait (Supplementary Table 13). Propiconazole concentration had a significant effect on siderophore (p = 0.0016), IAA (p = 0.0012), ammonia (p = 0.0138), and cellulase (p = 0.0012), but not on biofilm (p = 0.8488) or amylase (p = 0.9930). For siderophore and ammonia, only the highest dose (6.36 mg/L) differed significantly from all other treatments. IAA production at 0 mg/L differed significantly from all three propiconazole doses, which did not differ from one another. For cellulase, 0 mg/L and 2.12 mg/L did not differ from each other, nor did 4.24 mg/L and 6.36 mg/L, but both lower concentrations differed significantly from both higher concentrations. These results agree with the Wilcoxon signed-rank tests (Section 3.7.2).

##### 3.7.3.2 Multivariate PERMANOVA

Multivariate PERMANOVA on the six standardized traits (Euclidean distance, 999 permutations) showed a significant effect of propiconazole concentration on the overall PGP profile of the isolates (pseudo-F = 2.47, p = 0.001; Supplementary Table 14). The effect was small, however: concentration explained 6.0% of the total multivariate variation (R² = 0.060). Pairwise comparisons with FDR correction showed that the profiles at 0 mg/L differed from those at 4.24 mg/L (adjusted p = 0.014) and 6.36 mg/L (adjusted p = 0.012), and the profiles at 2.12 mg/L differed from those at 6.36 mg/L (adjusted p = 0.012). The two lowest concentrations did not differ from each other (adjusted p = 0.623), nor did 2.12 and 4.24 mg/L (adjusted p = 0.215) or 4.24 and 6.36 mg/L (adjusted p = 0.068). This agrees with the isolate-blocked PERMANOVA on the principal component scores (Section 3.7.6).

#### 3.7.4 PERMDISP Test

To determine whether groups differed in their dispersion around the centroid, PERMDISP was performed on standardized traits data. Global PERMDISP F = 2.06, p = 0.121 indicated that group dispersions were homogeneous (Supplementary Table 15). It can thus be assumed that the significant multivariate PERMANOVA result reflects differences in the position (centroids) of the treatment groups rather than differences in their spread (dispersion). Pesticide concentration groups had average dispersion ranging from 2.03 to 2.58, further consolidating the global PERMDISP outcome.

#### 3.7.5 Trait-wise Statistical Comparison

Trait-wise statistical comparison (Supplementary Table 16) showed using the IQR (Interquartile Range) rule that IAA and cellulase production had no outlier isolates. Isolates 24 and 26 were consistently flagged for high siderophore production. The large number of isolates flagged for biofilm and amylase mainly reflects the discrete biofilm scores and the many isolates without amylase activity, which compress the interquartile range, rather than true outliers. Trait-wise comparisons across concentrations are shown in Supplementary Figure 1.

#### 3.7.6 Principal component analysis (PCA)

PCA was performed to reduce dimensionality and identify the major axes of variation among the six PGPR traits. Six principal components were considered for the analysis, and it was observed that the first three principal components (PC1, PC2, and PC3) explained 66.75% of the total variance (Supplementary Table 17). Trait loadings show that biofilm and IAA contribute strongly to PC1, and siderophore moderatley, while ammonia has a weak positive contribution. Amylase and cellulase, on the other hand, are moderate negative contributors for PC1. For PC2, siderophore, amylase, and cellulase become strong positive contributors while IAA plays a moderately positive role and biofilm along with ammonia shows negligible effect. PC1 can be interpreted as the growth promotion-related axis while PC2 can be considered the enzymatic activity-related axis. PC3 was predominantly associated with ammonia and may therefore be interpreted as an ammonia-associated axis. The PCA centroids showed that concentration-related differences were primarily reflected along PC2 and PC3 (Supplementary Table 18; Supplementary Figure 2). PC2 scores shifted from positive values at 0 and 2.12 mg/L (0.377 and 0.184, respectively) to negative values at 4.24 and 6.36 mg/L (−0.192 and −0.369). Similarly, PC3 scores decreased from 0.363 and 0.279 at 0 and 2.12 mg/L to −0.111 and −0.531 at 4.24 and 6.36 mg/L, respectively. Both declines were significant in Friedman repeated-measures tests (PC2: χ² = 28.12, p < 0.001; PC3: χ² = 14.68, p = 0.002) and showed significant negative linear trends with concentration (PC2: r = −0.25, p = 0.006; PC3: r = −0.34, p < 0.001). PC1 scores also differed among concentrations (Friedman χ² = 10.04, p = 0.018) but showed no linear trend (r = 0.03, p = 0.755); they peaked at 4.24 mg/L (0.281) and were similar at the other concentrations (−0.125 to −0.035). An isolate-blocked PERMANOVA on PC1–PC3 scores confirmed a significant but small overall concentration effect (pseudo-F = 2.47, R² = 0.060, p = 0.0001). An interesting finding here is that siderophore loads positively on both PC1 (0.48) and PC2 (0.61).

#### 3.7.7 Redundancy Analysis (RDA)

Redundancy analysis (Supplementary Table 19) helped show that only 3.88% of the total variation can be explained by the pesticide concentrations (Observed R² = 0.0388), nd the association was statistically significant (permutation p = 0.001). Trait loadings show that biofilm showed a negligible positive association with concentration, whereas the other traits decreased.

#### 3.7.8 Ordinal Logistic Regression

Biofilm production was predicted by ordinal logistic regression (Supplementary Table 20). Siderophore production was the only significant predictor of biofilm category (β = 0.38, odds ratio = 1.46, p = 0.048).

#### 3.7.9 Kruskal–Wallis Test

The Kruskal–Wallis test was performed for non-parametric comparison of the treatment groups for each PGP trait, followed by Dunn’s post-hoc test with Benjamini–Hochberg FDR correction (Supplementary Table 21). Significant global differences were observed for ammonia (p = 0.0008), cellulase (p = 0.0035), and siderophore (p = 0.0499), but not for IAA, biofilm, or amylase. In pairwise comparisons, the highest propiconazole dose (6.36 mg/L) differed from all other treatments for ammonia (FDR-adjusted p = 0.0015–0.0060), and both lower concentrations (0 and 2.12 mg/L) differed from both higher concentrations (4.24 and 6.36 mg/L) for cellulase (FDR-adjusted p = 0.015–0.036). No siderophore comparison remained significant after FDR correction (lowest adjusted p = 0.057). Because the Kruskal–Wallis test treats the four concentration groups as independent samples, it does not account for the repeated measurement of the same isolates and is therefore less sensitive than the isolate-blocked test.

### 3.8 16S rDNA PCR and Sequencing

Genomic DNAs of all 30 isolates were extracted using Monarch BioLAB DNA extraction kit. Afterwards, 16S PCR was conducted using the following primer pair (Tables 8 and 9).

**Table 8:**
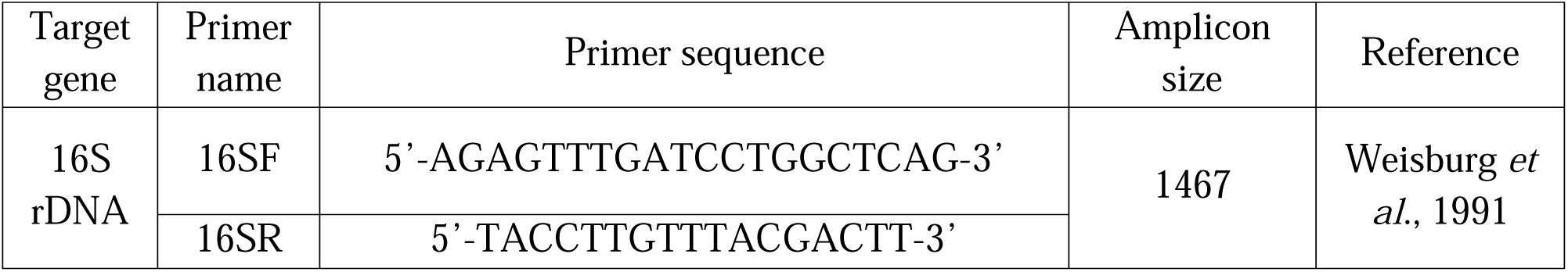
16S Primer names and sequences.

**Table 9:**
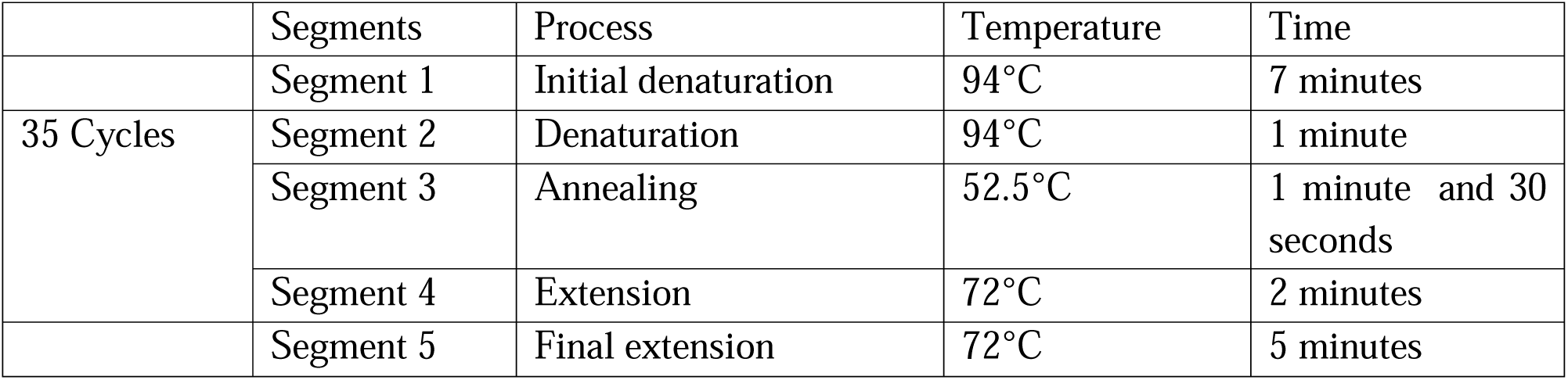
PCR reaction parameters.

After performing the PCR reaction, the PCR products were sequenced for identification. Twenty seven isolates were sequenced successfully, and they were identified by NCBI Nucleotide BLAST. *Bacillus* and *Agrobacterium* both were the most common genera, with six isolates each. Four *Priestia* and four *Staphylococcus* were found within the sequenced isolates, two were *Peribacillus huizhouensis -* and *Rhizobium sp., Faucicola osloensis, Lysinibacillus fusiformis, Sinorhizobium fredii,* and *Ensifer adhaerens* were each represented by one isolate (Table 10).

**Table 10:** Isolates and their accession numbers.

| Isolate | Accession No. and Taxonomy |
| --- | --- |
| 22 | PX052076.1 *Bacillus altitudinis* strain HAT20622 16S ribosomal RNA gene, partial sequence |
| 23 | ----- |
| 24 | PX052077.1 *Rhizobium* sp. strain HAT20624 16S ribosomal RNA gene, partial sequence |
| 26 | PX052328.1 *Agrobacterium* sp. strain HAT20626 16S ribosomal RNA gene, partial sequence |
| 27 | PX052329.1 *Agrobacterium tumefaciens* strain HAT20627 16S ribosomal RNA gene, partial sequence |
| 29 | PX052331.1 *Lysinibacillus fusiformis* strain HAT20629 16S ribosomal RNA gene, partial sequence |
| 30 | PX052332.1 *Priestia aryabhatai* strain HAT20630 16S ribosomal RNA gene, partial sequence |
| 34 | PX052333.1 *Priestia aryabhatai* strain HAT20634 16S ribosomal RNA gene, partial sequence |
| 35 | PX052334.1 *Sinorhizobium fredii* strain HAT20635 16S ribosomal RNA gene, partial sequence |
| 36 | PX052335.1 *Priestia megaterium* strain HAT20636 16S ribosomal RNA gene, partial sequence |
| 38 | PX052336.1 *Peribacillus huizhouensis* strain HAT20638 16S ribosomal RNA gene, partial sequence |
| 39 | PX052337.1 *Bacillus* sp. (in: firmicutes) strain HAT20639 16S ribosomal RNA gene, partial sequence |
| 40 | PX052338.1 *Bacillus* sp. (in: firmicutes) strain HAT20640 16S ribosomal RNA gene, partial sequence |
| 42 | PX052339.1 *Ensifer adhaerens* strain HAT20642 16S ribosomal RNA gene, partial sequence |
| 44 | PX052340.1 *Staphylococcus* sp. strain HAT20644 16S ribosomal RNA gene, partial sequence |
| 45 | PX052341.1 *Bacillus licheniformis* strain HAT20645 16S ribosomal RNA gene, partial sequence |
| 46 | PX052342.1 *Priestia* sp. strain HAT20646 16S ribosomal RNA gene, partial sequence |
| 47 | PX052343.1 *Peribacillus huizhouensis* strain HAT20647 16S ribosomal RNA gene, partial sequence |
| 48 | PX052344.1 *Agrobacterium salinitolerans* strain HAT20648 16S ribosomal RNA gene, partial sequence |
| 52 | PX052348.1 *Staphylococcus* sp. strain HAT20652 16S ribosomal RNA gene, partial sequence |
| 53 | PX052349.1 *Agrobacterium radiobacter* strain HAT20653 16S ribosomal RNA gene, partial sequence |
| 54 | PX052350.1 *Staphylococcus* sp. strain HAT20654 16S ribosomal RNA gene, partial sequence |
| 55 | PX052078.1 *Faucicola osloensis* strain HAT20655 16S ribosomal RNA gene, partial sequence |
| 56 | PX052351.1 *Staphylococcus* sp. strain HAT20656 16S ribosomal RNA gene, partial sequence |
| 57 | PX052352.1 *Agrobacterium* sp. strain HAT20657 16S ribosomal RNA gene, partial sequence |
| 58 | ----- |
| 59 | PX052353.1 *Agrobacterium pusense* strain HAT20659 16S ribosomal RNA gene, partial sequence |
| 60 | ----- |
| 63 | PX052355.1 *Bacillus* sp. (in: firmicutes) strain HAT20663 16S ribosomal RNA gene, partial sequence |
| 65 | PX052356.1 *Bacillus cereus* strain HAT20665 16S ribosomal RNA gene, partial sequence |

### 3.9 Scanning Electron Microscopy (SEM)

Isolate 53 (*Agrobacterium radiobacter* strain HAT20653) was selected as a representative organism to observe the effects of all four treatments of propiconazole on bacterial cell surface (Figure 1). Signs of cell surface damage were observed when the cells were challenged with propiconazole.

**Figure 1:**
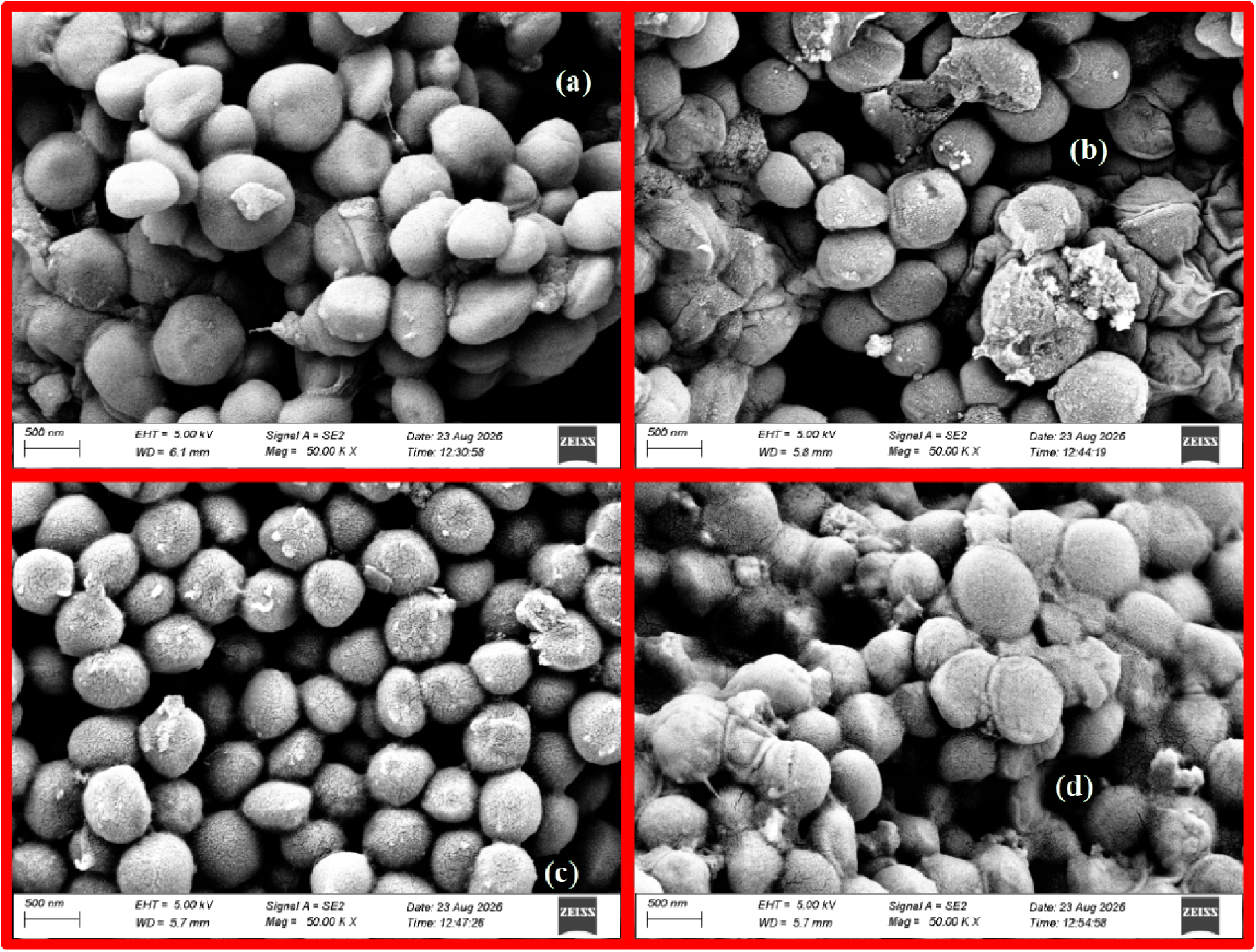
SEM images of *Agrobacterium radiobacter* strain HAT20653 under (a) 0 mg/L, (b) 2.12 mg/L, (c) 4.24 mg/L, and (d) 6.36 mg/L propiconazole treatment

## 4 Discussion

Propiconazole has been reported to increase microbial growth at lower concentrations but to inhibit it at higher concentration (Satapute *et al*., 2019). The fungicide has also shown the capacity of lowering soil dehydrogenase activity (Sliti *et al.,* 2024) and bacterial denitrification (Michel *et al.,* 2021). To further investigate this pesticide’s effects on rhizosphere bacteria, the current study was conducted using three propiconazole concentrations. The study intentionally used high concentrations of propiconazole to observe whether the pesticide has any off-target effect even at higher-than-normal levels. Our rationale was that if significant changes were observed at higher concentrations, the dose would be progressively reduced and the experiments repeated until the highest concentration that did not produce significant off-target effects was identified. Multivariate PERMANOVA revealed that the overall plant growth-promoting (PGP) profiles of the isolates changed significantly with increasing propiconazole concentrations; however, the effect size was small, with concentration accounting for only approximately 6% of the total variation. Similarly, redundancy analysis (RDA) showed that fungicide concentration explained only 3.9% of the variation in PGP profiles, although this effect was statistically significant. These findings were further supported by PERMDISP analysis, which indicated no significant differences in multivariate dispersion among treatment groups. As the lowest tested propiconazole concentration (2.12 mg/L) did not significantly alter the overall PGP profile, subsequent experiments were restricted to the tested concentrations.

Among the six PGP traits evaluated, biofilm formation was the least affected by propiconazole exposure. None of the statistical analyses detected significant differences among the four treatment groups, suggesting that bacterial biofilm formation in the rhizosphere is largely resilient to propiconazole stress. Interestingly, isolate 53, selected as the representative organism for SEM analysis, exhibited signs of cell surface disruption at all tested propiconazole concentrations, yet retained its strong biofilm-forming capacity across treatments. This observation suggests that while propiconazole exposure may induce structural alterations to the cell surface, the level of stress imposed is insufficient to substantially impair the adhesive properties required for biofilm formation.

Like biofilm formation, siderophore production data were analyzed using the Wilcoxon signed-rank test. Mean siderophore production at the highest propiconazole concentration (6.36 mg/L) was notably lower than that observed under the other treatments. This pattern was confirmed by the isolate-blocked permutation test, which showed that the 6.36 mg/L treatment differed significantly from all other concentrations. Although the Kruskal-Wallis test detected a significant overall difference among treatments (p = 0.0499), no pairwise comparison remained significant after FDR correction. This discrepancy is likely attributable to the inability of the Kruskal-Wallis test to account for repeated measurements of the same isolates. Collectively, these results suggest that siderophore production is adversely affected only at the highest propiconazole concentration, whereas lower concentrations have minimal impact.

In contrast, IAA production exhibited a different response pattern. Both the Wilcoxon signed-rank test and the isolate-blocked permutation test showed that the control treatment (0 mg/L) had significantly higher IAA production than all three propiconazole-treated groups, while no significant differences were detected among the three fungicide concentrations. The Kruskal-Wallis test, however, did not identify a significant overall treatment effect. Nevertheless, the consistent results obtained from analyses that accounted for isolate-specific variation indicate that exposure to propiconazole reduces IAA production. As no further decline was observed with increasing fungicide concentration, the inhibitory effect appears to occur upon exposure and does not intensify in a dose-dependent manner within the tested concentration range.

Mean ammonia production was significantly lowered under 6.36 mg/L propiconazole treatment, and this was also supported by the isolate-blocked permutation test along with the Kruskal– Wallis results. Research on the effect of propiconazole on bacterial ammonia production is scarce. The study by Satapute *et al*. (2019) states a lowered urease activity in soil due to propiconazole. This may partly explain the result in the current study as ammonia is released due to urease activity - but this alone should not be considered the only explanation as ammonia inside cell can be formed by numerous other reactions too.

Like biofilm formation, amylase production was not significantly affected by propiconazole exposure, suggesting that starch-degrading activity remains largely unaffected within the tested concentration range. In contrast, cellulase production exhibited a clear response to the fungicide. The Wilcoxon signed-rank test indicated that cellulase activity was comparable between the 0 and 2.12 mg/L treatments, as well as between the 4.24 and 6.36 mg/L treatments. The isolate-blocked permutation test produced the same grouping pattern, with both lower concentrations differing significantly from both higher concentrations following FDR correction. Consistent with these findings, the Kruskal-Wallis test detected a significant overall treatment effect (p = 0.0035), and Dunn’s post hoc test confirmed significant differences between the lower and higher concentration groups (FDR-adjusted p = 0.015-0.036). Collectively, these results indicate that cellulase production remains stable at lower propiconazole concentrations but declines at concentrations of 4.24 mg/L and above.

The PCA results suggest that the six PGPR traits can be broadly represented by three major patterns of variation. The concentration-related centroid patterns further indicate that increasing concentration affected these major patterns differently. PC1 scores did not show a consistent trend with concentration: they peaked at 4.24 mg/L but showed no linear relationship with dose, suggesting that the growth promotion-related axis did not display a clear concentration-dependent shift. In contrast, PC2 and PC3 showed distinct variations across concentrations, with their centroid scores generally shifting toward lower values at higher concentrations. This trend suggests that enzymatic activity and ammonia-related characteristics were more responsive to changes in pesticide concentration. This interpretation was further supported by the significant Friedman tests and negative linear trends for PC2 and PC3, whereas PC1, although significant in the Friedman test, showed no linear trend with concentration. Among the six PGP traits, siderophore contributed substantially to both of the first two principal components. This suggests that siderophore production may represent a functional link between the growth-promotion-related characteristics represented by PC1 and the enzyme-associated characteristics represented by PC2.

Redundancy analysis further demonstrated a significant association between pesticide concentration and the overall PGPR trait profile, although the proportion of variation explained by concentration was relatively small. The observed R² of 0.0388 indicates that concentration accounted for approximately 3.88% of the total multivariate variation, while the permutation test indicated that this association was statistically significant (p = 0.001). Overall, these findings indicate that increasing pesticide concentration was associated with a relatively modest but statistically detectable shift in the combined PGPR functional profile, with the response appearing to vary among individual traits rather than occurring uniformly across all six characteristics.

Ordinal logistic regression suggested an association between biofilm production and siderophore formation. This is supported by several preexisting works. Environmental concentration and chemical form of iron have been shown to regulate biofilm production in various bacteria (Kolodkin-Gal *et al*., 2013; Banin *et al*., 2005; Musk *et al*., 2005; Reid *et al*., 2009), establishing a link between these two traits. This has been discussed in case of *pvdF* mutant *Pseudomonas aeruginosa* (Harrison & Buckling, 2009). It was seen that siderophore-deficient bacteria were also deficient in biofilm formation. Similar results were found in *Bacillus subtilis* (Rizzi *et al*., 2019), where a biofilm-deficient mutant (*epsA-O tasA*) showed severely impeded growth due to an inability to efficiently recover Fe-bacillibactin. Furthermore, another study on *Staphylococcus epidermidis* strengthened the idea with the finding that siderophore biosynthesis was critical for biofilm formation under iron-restricted conditions where Δsfa isolates formed sparse biofilms (Oliveira *et al*., 2021), indicating *sfa ABCD* having a crucial role in biofilm formation. According to Oh *et al*. (2013), the *rpoS* sigma factor negatively regulates production of IAA and siderophore in a biocontrol rhizobacterium, *Pseudomonas chlororaphis* O6. On the other hand, the *TonB* system has been shown to functionally link siderophore-mediated iron uptake with cellulose utilization in *Teredinibacter turnerae*, as *tonB1b* and *tonB2* are required for both processes (Naka and Haygood, 2023). Moreover, maltose, an intermediate product of starch degradation, was reported to induce the biosynthesis of the siderophore lysochelin in *Lysobacter* sp. 3655 in rich media that are not compatible with siderophore production (Zhang *et al.,* 2024) - pointing to a possible interplay between amylase and siderophore. All these references together suggest siderophore may act as a bridging link between the growth promotion related axis and the enzymatic activity associated axis as perceived in the PCA results, although the exact mechanism remains largely unknown.

The present study suggests that propiconazole affects wheat rhizosphere bacteria in a selective, trait-specific manner rather than causing broad disruption of PGPR functions. Although the overall PGP profile changed significantly with increasing propiconazole concentration, the magnitude of this effect was small, indicating that only certain traits were affected. However, these findings should be validated under soil and plant growth conditions. The study also has several limitations. Because the bacterial isolates belonged to multiple species, their metabolic diversity prevented detailed investigation of the molecular mechanisms underlying changes in individual PGP traits. In addition, the effect of propiconazole on cell surface structure was examined in only one representative isolate. Therefore, these results should be considered preliminary and serve as a basis for future studies aimed at understanding the specific mechanisms through which propiconazole influences PGPR functionality.

## Data availability statement

The 16S rDNA genetic data of 27 isolates were sequenced, characterized, and deposited into NCBI database under the accession numbers PX052076-PX052078, PX052328-PX052329, PX052331-PX052344, PX052348-PX052353, and PX052355-PX052356.

## Author Contributions

MTH - Conceptualization, Data curation, Formal Analysis, Investigation, Methodology, Visualization, Writing – original draft.

AS - Data curation, Formal Analysis, Investigation, Methodology, Writing – original draft.

HA - Investigation, Methodology, Supervision, Validation, Funding acquisition, Project administration, Resources, Writing – review & editing.

AB - Validation, Project administration, Resources, Writing – review & editing.

## Acknowledgement of Funding

The author(s) gratefully acknowledge that this work was supported by the Ministry of Science and Technology, Government of the People’s Republic of Bangladesh. The funding was awarded to the following project: “Mechanism and application of plant growth-promoting rhizobacteria: an alternative to chemical fertilizers for sustainable agricultural development.” Project ID: SRG-233514.

## Ethics and Consent to Participate declarations

Not applicable

## Generative AI statement

During the preparation of this manuscript, the authors used Claude (Anthropic), an AI assistant, to help write and check statistical analysis scripts, prepare tables, and check the consistency of the manuscript with the Supplementary Material. The authors reviewed and edited all AI-assisted content and take full responsibility for the content of this publication.

