## Supplementary Material for "Study on The Effect of a Propiconazole Based Fungicide on Plant Growth Promoting Properties of Isolates from Wheat Rhizosphere"

**Supplementary Materials**

**Normality Test**

**Table 1: Statistical Assessment of Normality for Siderophore Data Using Shapiro–Wilk and Kolmogorov–Smirnov Tests**

| Treatment | N | Shapiro-Wilk p | KS p-value | Normal? |
| --- | --- | --- | --- | --- |
| 0 mg/L | 30 | **0.000358** | 0.337688 | No |
| 2.12 mg/L | 30 | **0.000724** | 0.390983 | No |
| 4.24 mg/L | 30 | **0.001175** | 0.454591 | No |
| 6.36 mg/L | 30 | **1e-06** | **0.031949** | No |

**Table 2: Statistical Assessment of Normality for IAA Data Using Shapiro–Wilk and Kolmogorov–Smirnov Tests**

| Treatment | N | Shapiro-Wilk p | KS p-value | Normal? |
| --- | --- | --- | --- | --- |
| 0 mg/L | 30 | **0.000503** | **0.026233** | No |
| 2.12 mg/L | 30 | **9.3e-05** | **0.023767** | No |
| 4.24 mg/L | 30 | **5.3e-05** | **0.014315** | No |
| 6.36 mg/L | 30 | **9e-05** | 0.060769 | No |

**Table 3: Statistical Assessment of Normality for Ammonia Data Using Shapiro–Wilk and Kolmogorov–Smirnov Tests**

| Treatment | N | Shapiro-Wilk p | KS p-value | Normal? |
| --- | --- | --- | --- | --- |
| 0 mg/L | 30 | 0.661693 | 0.758495 | Yes |
| 2.12 mg/L | 30 | 0.23174 | 0.552566 | Yes |
| 4.24 mg/L | 30 | 0.356269 | 0.904991 | Yes |
| 6.36 mg/L | 30 | **0.001115** | 0.567253 | No |

**Table 4: Statistical Assessment of Normality for Biofilm Data Using Shapiro–Wilk and Kolmogorov–Smirnov Tests**

| Treatment | N | Shapiro-Wilk p | KS p-value | Normal? |
| --- | --- | --- | --- | --- |
| 0 mg/L | 30 | **6e-06** | **0.000148** | No |
| 2.12 mg/L | 30 | **5.3e-05** | **0.001671** | No |
| 4.24 mg/L | 30 | **3.6e-05** | **0.000489** | No |
| 6.36 mg/L | 30 | **1.8e-05** | **0.000318** | No |

**Table 5: Statistical Assessment of Normality for Cellulase Data Using Shapiro–Wilk and Kolmogorov–Smirnov Tests**

| Treatment | N | Shapiro-Wilk p | KS p-value | Normal? |
| --- | --- | --- | --- | --- |
| 0 mg/L | 30 | **0.000807** | **0.022981** | No |
| 2.12 mg/L | 30 | 0.253306 | 0.893885 | Yes |
| 4.24 mg/L | 30 | **2e-05** | **0.024419** | No |
| 6.36 mg/L | 30 | **0.002771** | 0.13843 | No |

**Table 6: Statistical Assessment of Normality for Amylase Data Using Shapiro–Wilk and Kolmogorov–Smirnov Tests**

| Treatment | N | Shapiro-Wilk p | KS p-value | Normal? |
| --- | --- | --- | --- | --- |
| 0 mg/L | 30 | **0.0** | **0.002185** | No |
| 2.12 mg/L | 30 | **0.0** | **0.000727** | No |
| 4.24 mg/L | 30 | **0.0** | **1e-06** | No |
| 6.36 mg/L | 30 | **0.0** | **1.2e-05** | No |

**Wilcoxon Test**

**Table 7: Pairwise Wilcoxon Signed‑Rank Test Results for Siderophore Production Under Different Pesticide Concentrations. Values Represent p‑values with Significance Codes (*p < 0.05, **p < 0.01, ***p < 0.001, ns = not significant)**

| Treatment | 0 mg/L | 2.12 mg/L | 4.24 mg/L | 6.36 mg/L |
| --- | --- | --- | --- | --- |
| 0 mg/L | — | 0.0544 ns | 0.2301 ns | 0.0005 *** |
| 2.12 mg/L | 0.0544 ns | — | 0.9757 ns | 0.0007 *** |
| 4.24 mg/L | 0.2301 ns | 0.9757 ns | — | 0.0068 ** |
| 6.36 mg/L | 0.0005 *** | 0.0007 *** | 0.0068 ** | — |

**Table 8: Pairwise Wilcoxon Signed‑Rank Test Results for IAA Production Under Different Pesticide Concentrations. Values Represent p‑values with Significance Codes (*p < 0.05, **p < 0.01, ***p < 0.001, ns = not significant)**

| Treatment | 0 mg/L | 2.12 mg/L | 4.24 mg/L | 6.36 mg/L |
| --- | --- | --- | --- | --- |
| 0 mg/L | — | 0.0000 *** | 0.0015 ** | 0.0009 *** |
| 2.12 mg/L | 0.0000 *** | — | 0.3184 ns | 0.5237 ns |
| 4.24 mg/L | 0.0015 ** | 0.3184 ns | — | 0.0880 ns |
| 6.36 mg/L | 0.0009 *** | 0.5237 ns | 0.0880 ns | — |

**Table 9: Pairwise Wilcoxon Signed‑Rank Test Results for Ammonia Production Under Different Pesticide Concentrations. Values Represent p‑values with Significance Codes (*p < 0.05, **p < 0.01, ***p < 0.001, ns = not significant)**

| Treatment | 0 mg/L | 2.12 mg/L | 4.24 mg/L | 6.36 mg/L |
| --- | --- | --- | --- | --- |
| 0 mg/L | — | 0.5425 ns | 1.0000 ns | 0.0047 ** |
| 2.12 mg/L | 0.5425 ns | — | 0.5425 ns | 0.0015 ** |
| 4.24 mg/L | 1.0000 ns | 0.5425 ns | — | 0.0006 *** |
| 6.36 mg/L | 0.0047 ** | 0.0015 ** | 0.0006 *** | — |

**Table 10: Pairwise Wilcoxon Signed‑Rank Test Results for Biofilm Production Under Different Pesticide Concentrations. Values Represent p‑values with Significance Codes (*p < 0.05, **p < 0.01, ***p < 0.001, ns = not significant)**

| Treatment | 0 mg/L | 2.12 mg/L | 4.24 mg/L | 6.36 mg/L |
| --- | --- | --- | --- | --- |
| 0 mg/L | — | 0.7815 ns | 0.2720 ns | 0.4174 ns |
| 2.12 mg/L | 0.7815 ns | — | 0.1303 ns | 0.2180 ns |
| 4.24 mg/L | 0.2720 ns | 0.1303 ns | — | 0.9141 ns |
| 6.36 mg/L | 0.4174 ns | 0.2180 ns | 0.9141 ns | — |

**Table 11: Pairwise Wilcoxon Signed‑Rank Test Results for Cellulase Production Under Different Pesticide Concentrations. Values Represent p‑values with Significance Codes (*p < 0.05, **p < 0.01, ***p < 0.001, ns = not significant)**

| Treatment | 0 mg/L | 2.12 mg/L | 4.24 mg/L | 6.36 mg/L |
| --- | --- | --- | --- | --- |
| 0 mg/L | — | 0.0619 ns | 0.0007 *** | 0.0021 ** |
| 2.12 mg/L | 0.0619 ns | — | 0.0001 *** | 0.0036 ** |
| 4.24 mg/L | 0.0007 *** | 0.0001 *** | — | 0.1361 ns |
| 6.36 mg/L | 0.0021 ** | 0.0036 ** | 0.1361 ns | — |

**Table 12: Pairwise Wilcoxon Signed‑Rank Test Results for Amylase Production Under Different Pesticide Concentrations. Values Represent p‑values with Significance Codes (*p < 0.05, **p < 0.01, ***p < 0.001, ns = not significant)**

| Treatment | 0 mg/L | 2.12 mg/L | 4.24 mg/L | 6.36 mg/L |
| --- | --- | --- | --- | --- |
| 0 mg/L | — | 0.6273 ns | 0.0700 ns | 0.2667 ns |
| 2.12 mg/L | 0.6273 ns | — | 0.2719 ns | 0.5509 ns |
| 4.24 mg/L | 0.0700 ns | 0.2719 ns | — | 0.6460 ns |
| 6.36 mg/L | 0.2667 ns | 0.5509 ns | 0.6460 ns | — |

**PERMANOVA**

**Table 13: Isolate‑Blocked Permutation Test Results for Plant Growth‑Promoting Traits Under Different Pesticide Concentrations. Values Represent p‑values with Significance Codes (*p < 0.05, **p < 0.01, ***p < 0.001, ns = not significant)**

| Trait | Global test (Holm‑adjusted p) | Pairwise comparisons, mg/L (BH‑FDR‑adjusted p) | | | | | |
| --- | --- | --- | --- | --- | --- | --- | --- |
|  |  | 0 vs 2.12 | 0 vs 4.24 | 0 vs 6.36 | 2.12 vs 4.24 | 2.12 vs 6.36 | 4.24 vs 6.36 |
| Siderophore | 0.0016 ** | 0.1482 ns | 0.3660 ns | 0.0012 ** | 0.8660 ns | 0.0018 ** | 0.0192 * |
| IAA | 0.0012 ** | 0.0012 ** | 0.0144 * | 0.0344 * | 0.8126 ns | 0.3828 ns | 0.0750 ns |
| Ammonia | 0.0138 * | 0.7745 ns | 0.9288 ns | 0.0428 * | 0.7560 ns | 0.0108 * | 0.0234 * |
| Biofilm | 0.8488 ns | 1.0000 ns | 0.7836 ns | 0.8031 ns | 0.7836 ns | 0.7836 ns | 1.0000 ns |
| Cellulase | 0.0012 ** | 0.0749 ns | 0.0006 *** | 0.0021 ** | 0.0006 *** | 0.0012 ** | 0.0854 ns |
| Amylase | 0.9930 ns | 0.9928 ns | 0.9928 ns | 0.9928 ns | 0.9928 ns | 0.9928 ns | 0.9928 ns |

**Table 14: Multivariate PERMANOVA Results(Global and Pairwise, Distance: Euclidean. PERMANOVA with 999 permutations. Pairwise p-values adjusted using Benjamini–Hochberg FDR)**

| Global PERMANOVA | | | | | | | | | | | | | |
| --- | --- | --- | --- | --- | --- | --- | --- | --- | --- | --- | --- | --- | --- |
| method name | | test statistic name | | sample size | number of groups | test statistic | | p-value | number of permutations | | R² | | |
| PERMANOVA | | pseudo-F | | 120 | 4 | 2.4673712879639877 | | 0.001 | 999 | | 0.059983687735335106 | | |
| Pairwise PERMANOVA (with FDR correction) | | | | | | | | | | | | | |
| Group1 | Group2 | | pseudo-F | | | | R² | | | p-value | | Adj p-value (FDR) | Significant |
| 0 mg/L | 2.12 mg/L | | 0.712189750618116 | | | | 0.012130185463072736 | | | 0.623 | | 0.623 | No |
| 0 mg/L | 4.24 mg/L | | 3.004116805646161 | | | | 0.049244493043264885 | | | 0.007 | | 0.014 | Yes |
| 0 mg/L | 6.36 mg/L | | 3.588508026728555 | | | | 0.058265870398600865 | | | 0.004 | | 0.012 | Yes |
| 2.12 mg/L | 4.24 mg/L | | 1.5005104037324544 | | | | 0.02521844591837867 | | | 0.179 | | 0.2148 | No |
| 2.12 mg/L | 6.36 mg/L | | 3.8385788184443177 | | | | 0.06207417589130302 | | | 0.003 | | 0.012 | Yes |
| 4.24 mg/L | 6.36 mg/L | | 2.213177728474284 | | | | 0.036755703850316665 | | | 0.045 | | 0.0675 | No |

### Table 15: PERMDISP Results

| F-statistic | p-value | | Permutations |
| --- | --- | --- | --- |
| 2.0597 | 0.1210 | | 999 |
| Group Dispersions | | | |
| Group | | Average Dispersion | |
| 0 mg/L | | 2.5765 | |
| 2.12 mg/L | | 2.1138 | |
| 4.24 mg/L | | 2.1086 | |
| 6.36 mg/L | | 2.0315 | |
| Ranked Groups by Dispersion (Most → Least) | | | |
| Group | | Average Dispersion | |
| 0 mg/L | | 2.5765 | |
| 2.12 mg/L | | 2.1138 | |
| 4.24 mg/L | | 2.1086 | |
| 6.36 mg/L | | 2.0315 | |

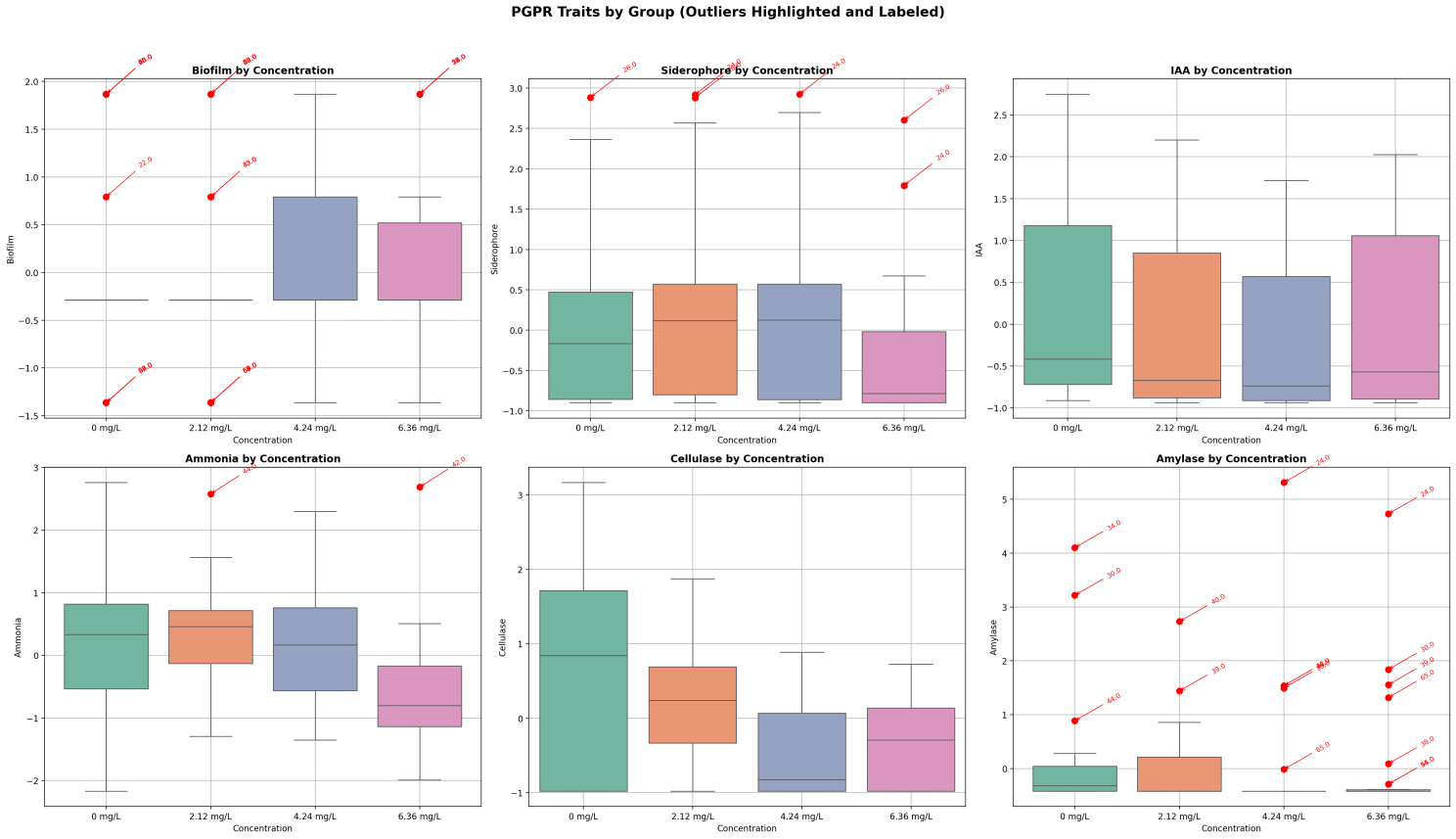

**Supplementary Figure 1: Trait-wise Statistical Comparison**

**Table 16: Outlier Isolates in Different PGP Properties Under Different Propiconazole Treatment**

| **Trait** | **0 mg/L** | **2.12 mg/L** | **4.24 mg/L** | **6.36 mg/L** |
| --- | --- | --- | --- | --- |
| Siderophore | **n = 1**  26 (2.883) | **n = 2**  24 (2.916)  26 (2.879) | **n = 1**  24 (2.924) | **n = 2**  24 (1.792)  26 (2.603) |
| IAA | — | — | — | — |
| Ammonia | — | **n = 1**  44 (2.578) | — | **n = 1**  42 (2.686) |
| Biofilm | **n = 11**  22 (0.790)  23 (−1.364)  26 (1.867)  48 (1.867)  52 (−1.364)  53 (1.867)  55 (1.867)  56 (−1.364)  57 (−1.364)  60 (−1.364)  63 (1.867) | **n = 14**  22 (1.867)  26 (1.867)  35 (0.790)  42 (0.790)  48 (1.867)  52 (−1.364)  53 (1.867)  54 (−1.364)  55 (1.867)  56 (−1.364)  58 (−1.364)  59 (−1.364)  60 (−1.364)  65 (−1.364) | — | **n = 6**  22 (1.867)  26 (1.867)  29 (1.867)  53 (1.867)  55 (1.867)  57 (1.867) |
| Cellulase | — | — | — | — |
| Amylase | **n = 3**  30 (3.218)  34 (4.102)  44 (0.887) | **n = 2**  39 (1.440)  40 (2.730) | **n = 5**  24 (5.311)  34 (1.538)  39 (1.494)  40 (1.538)  65 (−0.015) | **n = 7**  24 (4.730)  30 (1.840)  38 (0.088)  39 (1.556)  44 (−0.291)  56 (−0.291)  65 (1.319) |

**Table 17: Loadings (Trait–Component Correlations) of Plant Growth‑Promoting Traits on Principal Components PC1–PC6**

| Trait | PC1 | PC2 | PC3 | PC4 | PC5 | PC6 |
| --- | --- | --- | --- | --- | --- | --- |
| Siderophore | 0.484 | **0.615** | −0.224 | −0.033 | **0.566** | −0.129 |
| IAA | **0.683** | 0.324 | 0.098 | −0.413 | −0.448 | −0.219 |
| Ammonia | 0.293 | 0.022 | **0.896** | −0.010 | 0.173 | 0.283 |
| Biofilm | **0.707** | −0.016 | −0.167 | **0.634** | −0.201 | 0.173 |
| Cellulase | −0.453 | **0.576** | 0.352 | 0.385 | −0.134 | −0.416 |
| Amylase | −0.345 | **0.734** | −0.162 | −0.078 | −0.180 | **0.527** |
| Eigenvalue | 1.610 | 1.353 | 1.042 | 0.729 | 0.641 | 0.625 |
| Variance (%) | 26.84 | 22.55 | 17.36 | 12.14 | 10.69 | 10.42 |
| Cumulative (%) | 26.84 | 49.39 | 66.75 | 78.89 | 89.58 | 100.00 |

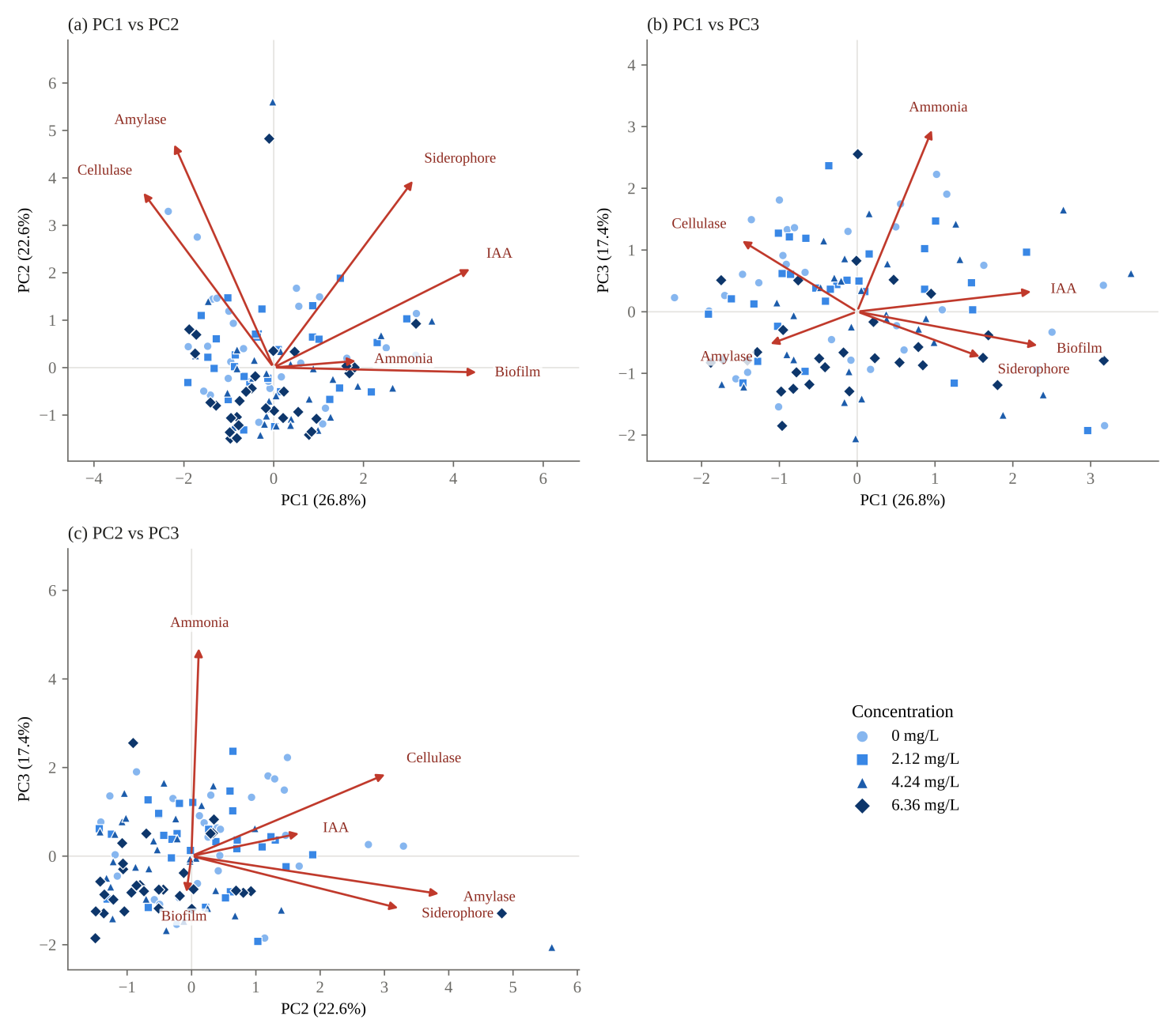
**Supplementary Figure 2: PCA Biplot**

**Table 18: Principal Component Analysis Summary: Variance Explained, Eigenvector Coefficients, Concentration Centroids and Tests of Pesticide‑Concentration Effects on PC1–PC3 Scores. Significance Codes (*p < 0.05, **p < 0.01, ***p < 0.001, ns = not significant)**

| **Parameter** | **PC1** | **PC2** | **PC3** |
| --- | --- | --- | --- |
| **Variance explained** | | | |
| Variance (%) | 26.84 | 22.55 | 17.36 |
| Cumulative variance (%) | 26.84 | 49.39 | 66.75 |
| **Eigenvector coefficients** | | | |
| Biofilm | 0.5568 | −0.0136 | −0.1637 |
| Siderophore | 0.3816 | 0.5283 | −0.2195 |
| IAA | 0.5381 | 0.2787 | 0.0965 |
| Ammonia | 0.2308 | 0.0190 | 0.8784 |
| Cellulase | −0.3571 | 0.4948 | 0.3447 |
| Amylase | −0.2722 | 0.6308 | −0.1591 |
| **Concentration centroids (mean PC score)** | | | |
| 0 mg/L | −0.1248 | 0.3766 | 0.3627 |
| 2.12 mg/L | −0.0349 | 0.1843 | 0.2785 |
| 4.24 mg/L | 0.2806 | −0.1918 | −0.1105 |
| 6.36 mg/L | −0.1209 | −0.3691 | −0.5307 |
| **Friedman test (df = 3)** | | | |
| χ² | 10.040 | 28.120 | 14.680 |
| p‑value | **0.0182 *** | **3.43 × 10^−6^ ***** | **0.0021 **** |
| **Linear trend across concentrations** | | | |
| Slope | 0.0154 | −0.1233 | −0.1448 |
| r | 0.0288 | −0.2512 | −0.3362 |
| R² | 0.0008 | 0.0631 | 0.1130 |
| p‑value | 0.7546 ns | **0.0057 **** | **0.0002 ***** |
| **Blocked PERMANOVA (PC1–PC3 jointly; isolate as block)** | | | |
| Result | **Pseudo‑F = 2.4674; R² = 0.0600; p = 0.0001 ***** | | |

### Table 19: Redundancy Analysis (RDA) Results

| Observed R² (variance explained): 0.0388 | | | | | | |
| --- | --- | --- | --- | --- | --- | --- |
| Permutation p-value: 0.0010 | | | | | | |
| Trait Loadings (Predictor Coefficients) | | | | | | |
| Predictor | Biofilm | Siderophore | IAA | Ammonia | Cellulase | Amylase |
| Concentration | 0.0646 | -0.1071 | -0.0670 | -0.2286 | -0.3370 | -0.0224 |

**Table 20: Biofilm Prediction Using Ordinal Logistic Regression**

Regression Equation:

Biofilm ~ -0.2258*Amylase + -0.2264*Cellulase + 0.3803*Siderophore + 0.0469*Ammonia + 0.2442*IAA

#### Detailed Results

| Predictor | Coefficient | Std. Error | z-value | p-value | 95% CI Lower | 95% CI Upper |
| --- | --- | --- | --- | --- | --- | --- |
| Amylase | -0.2258 | 0.182 | -1.241 | 0.2146 | -0.5825 | 0.1308 |
| Cellulase | -0.2264 | 0.1919 | -1.1798 | 0.2381 | -0.6024 | 0.1497 |
| Siderophore | 0.3803 | 0.192 | 1.981 | 0.0476 | 0.004 | 0.7567 |
| Ammonia | 0.0469 | 0.1789 | 0.2619 | 0.7934 | -0.3038 | 0.3975 |
| IAA | 0.2442 | 0.198 | 1.2332 | 0.2175 | -0.1439 | 0.6323 |
| -1.364120501373702/-0.2871832634470952 | -1.8453 | 0.2647 | -6.971 | 0.0 | -2.3641 | -1.3265 |
| -0.2871832634470952/0.789753974479512 | 1.0791 | 0.1049 | 10.287 | 0.0 | 0.8735 | 1.2847 |
| 0.789753974479512/1.866691212406119 | -0.6651 | 0.322 | -2.0657 | 0.0389 | -1.2962 | -0.034 |

**Table 21: Kruskal–Wallis Test Results with Dunn’s Post‑hoc Pairwise Comparisons for Plant Growth‑Promoting Traits Under Different Pesticide Concentrations. Values Represent p‑values with Significance Codes (*p < 0.05, **p < 0.01, ***p < 0.001, ns = not significant)**

| Trait | Kruskal–Wallis p‑value | Dunn’s pairwise comparisons, mg/L (BH‑FDR‑adjusted p) | | | | | |
| --- | --- | --- | --- | --- | --- | --- | --- |
|  |  | 0 vs 2.12 | 0 vs 4.24 | 0 vs 6.36 | 2.12 vs 4.24 | 2.12 vs 6.36 | 4.24 vs 6.36 |
| Siderophore | **0.0499 *** | 0.7534 ns | 0.7534 ns | 0.1262 ns | 0.9028 ns | 0.0573 ns | 0.0573 ns |
| IAA | 0.2442 ns | Not performed (global test not significant) | | | | | |
| Ammonia | **0.0008 ***** | 0.7694 ns | 0.7694 ns | **0.0034 **** | 0.7280 ns | **0.0015 **** | **0.0060 **** |
| Biofilm | 0.6737 ns | Not performed (global test not significant) | | | | | |
| Cellulase | **0.0035 **** | 0.9370 ns | **0.0145 *** | **0.0360 *** | **0.0145 *** | **0.0360 *** | 0.6901 ns |
| Amylase | 0.0540 ns | Not performed (global test not significant) | | | | | |
